# Stability-instability transition in tripartite merged ecological networks

**DOI:** 10.1101/2022.03.18.484866

**Authors:** Clive Emary, Anne-Kathleen Malchow

## Abstract

Although ecological networks are typically constructed based on a single type of interaction, e.g. trophic interactions in a food web, a more complete picture of ecosystem composition and functioning arises from merging networks of multiple interaction types. In this work, we consider tripartite networks constructed by merging two bipartite networks, one mutualistic and one antagonistic. Taking the interactions within each sub-network to be distributed randomly, we consider the stability of the dynamics of the network based on the spectrum of its community matrix. In the asymptotic limit of a large number of species, we show that the spectrum undergoes an eigenvalue phase transition, which leads to an abrupt destabilisation of the network as the ratio of mutualists to antagonists is increased. We also derive results that show how this transition is manifest in networks of finite size, as well as when disorder is introduced in the segregation of the two interaction types. Our random-matrix results will serve as a baseline for understanding the behaviour of merged networks with more realistic structures and/or more detailed dynamics.

## I. INTRODUCTION

A central goal of community ecology is to identify the mechanisms that maintain biodiversity in natural communities. One way to approach this problem is through the lens of ecological networks — abstract representations of interactions between taxa in a community [1–3]. Classically, studies of ecological networks have tended to focus on a single type of interaction, e.g. trophic interactions in a food web [4–6] or mutualistic interactions in a plant-pollinator network [7, 8]. Over the last decade or so, however, there has been a growing recognition that further progress requires the construction and study of networks that contain multiple interaction types [9–11] and even combining ecological with social interactions [12].

In this paper we consider the stability of ecological networks based on the tripartite structure of Fig. 1. These networks consist of three guilds (nominally plants, herbivores and mutualists) interacting through two distinct sets of interactions, one antagonistic and the other mutualistic. This tripartite structure represents a minimal example of a merged network [10, 13], and reflects the large-scale structure seen in empirical networks such as those discussed by Ref. 14–19. In the framework of multi-layer networks [20] Fig. 1 could be viewed as a two-layer multiplex network with antagonists in one layer, mutualists in the other and plants common to both. Two recent surveys analysed structural features and robustness of a number of tripartite networks from the literature, including those with mutualist and antagonist partitions as considered here; Ref. 21 found that the importance of species is positively correlated between the two bipartite subnetworks, and Ref. 22 found that the robustness (with respect to plant losses) of tripartite mutualist-antagonist networks could be understood in terms of the robustnesses of the two bipartite networks composing them.

**FIG. 1.**
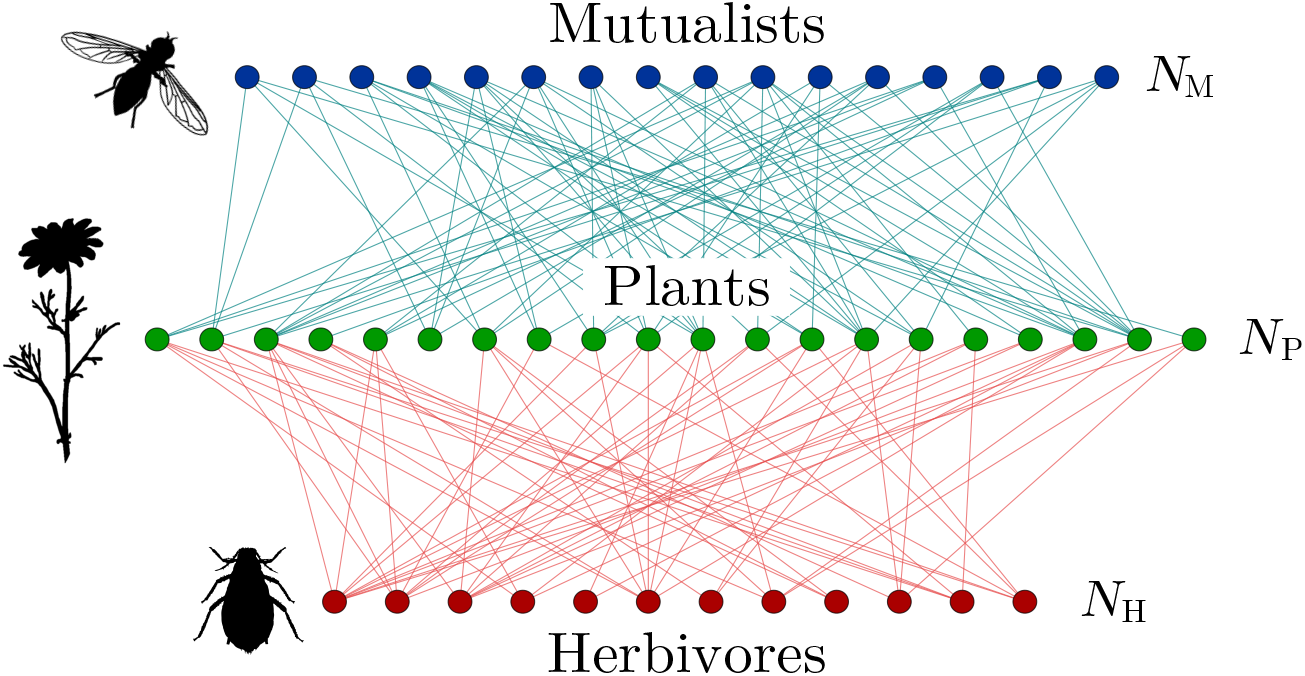
A tripartite ecological network consisting of *N*_H_ herbivores and *N*_M_ mutualists interacting with a set of *N*_P_ plants. Without disorder, all interactions between herbivores and plants are antagonistic (red edges), whereas all interactions between mutualists and plants are mutualistic (blue). Taxa images are public domain from http://www.phylopic.org.

The aim of the present contribution is to provide an analysis of the dynamical stability of this network structure when the subnetwork interactions are described by random matrices. The use of random matrices to shed light on ecosystem stability has a long history [23, 24] and acts as a baseline scenario against which more detailed and ecologically-motivated studies can be compared [25]. The stability of networks comprising a mixture of antagonistic and mututalistic interactions has previously been considered from a random-matrix viewpoint [26–28]. However, these studies have been of (stochastically) homogeneous models, i.e. without the tripartite structure of Fig. 1. The persistence and stability of an ecosystem with this structure was considered in Ref. 13 using numerical simulations for small networks. In contrast, the focus of our work is on analytic results for ecosystems consisting of a large number of species.

Specifically we investigate the stability of a community matrix with block-structure that reflects Fig. 1 and with random interactions within the blocks. Using the results of Ref. 29 and Ref. 30 we give an account of key spectral features of such matrices in the limit of large network size. We show that the spectrum consists of a bulk component and a pair of eigenvalues of large magnitude. The properties of these latter split the behaviour of the model into two distinct phases. In the antagonist-dominated phase, the large eigenvalues are complex and the stability properties of the model are determined by the bulk. In the mutualist-dominated phase, the large eigenvalues are real and serve to destabilise the system. The transition between these two phases represents an eigenvalue phase transition [31] and can be driven, for example, by an increase in the relative fraction of mutualists in the ecosystem. Whilst this phase transition strictly takes place in the asymptotic limit, we also derive an expression for the stability-determining eigenvalue valid at finite system size. Finally, we also consider a scenario in which we introduce a degree of disorder into the network of Fig. 1 by exchanging the type of a certain random fraction of interactions. We show that the phase transition can survive in the presence of disorder, but vanishes if the disorder is too strong.

## II. TRIPARTITE NETWORK MODEL

We consider a system of *N*_P_ plants (or producers), *N*_M_ mutualists and *N*_M_ herbivores such that the total number of consumers (mutualists and herbivores) is *N*_C_ = *N*_M_ + *N*_H_. We then define *s* = *N*_C_/*N*_P_ as the ratio of consumers to plants and *r* = *N*_M_/*N*_C_ as the ratio of mutualists to consumers. We assume that, close to equilibrium, the dynamics of the populations is described by community matrix *K* = −*D* + *A*, where *D* describes intraspecific competition and *A* describes interspecific interaction. As is conventional, we assume *D* to have elements *D_ij_* = *dδ_ij_* with *δ_ij_* the Kronecker delta symbol and *d* the competition strength. We then choose *K* to reflect the tripartite structure of the network in Fig. 1. In the first instance, we take *A* to have the following block structure

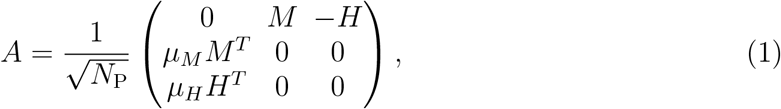

in which the species have been ordered plants first, then mutualists and finally herbivores. The block *M* is of dimension *N*_P_ × *N*_M_ and describes the plant-mutualist interactions; block *H* is of dimension *N*_P_ × *N*_H_ and describes plant-herbivore interactions. We take all matrix elements of *M* and *H* to be positive such that the the nature of the interactions is encoded in the explicit signs shown in Eq. (1). Parameters *μ_M_* and *μ_H_* are included to describe asymmetries in the two directions of each interaction type. The forefactor 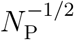 is included for mathematical convenience. We note that a similar matrix block structure was considered by Ref. 32 but with sign assignments appropriate to trophic interactions.

This overall structure is then supplemented by a random model for the matrix elements of the blocks. Considering the mutualists, we set 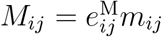 where the interaction coefficients *m_ij_* are treated as independent identical random variables with mean 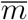 and variance 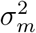, and where the factors 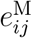 describe the links of the plant-mutualist sub-network: species that interact have 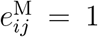; those that do not have 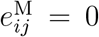. The mutualist subnetwork connectance is therefore 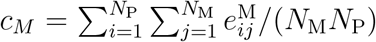. We assume a random structure for the plant-mutualist network with 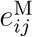 chosen from a Bernoulli distribution with probability 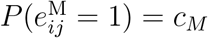 for all pairs *i,j*. The first and second cumulants of the matrix elements of *M* are thus

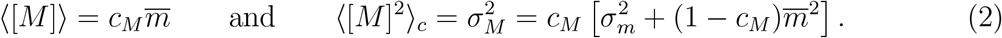

An analogous set-up and notation is employed for the matrix elements of *H*.

## III. EIGENVALUE SPECTRUM

The stability of the equilibrium described by community matrix *K* is determined by its *N*_C_ + *N*_P_ eigenvalues which we denote *ϵ*. Multiplying out the block structure of eigenvalue equation *K***v** = *ϵ***v**, we find that the non-trivial eigenvalues of *K* are given by Ref. 33

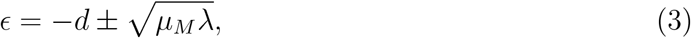

where *λ* are obtained from the eigenvalue equation

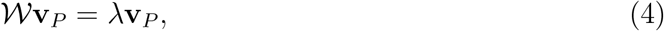

in which **v**_*P*_ is a length-*N*_P_ vector and

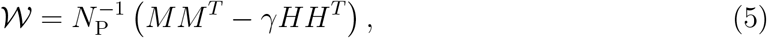

is an *N*_P_ × *N*_P_ matrix with *γ* = *μ_H_*/*μ_M_*. The stability of community matrix *K* therefore becomes a question of the spectrum of 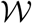. Since 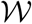 is symmetric, its eigenvalues are real and a value *λ* > 0 contributes two real eigenvalues to the spectrum of *K*; conversely, a value of *λ* < 0 contributes an imaginary pair. For *s* > 1, i.e. more consumers than plants, matrix 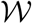 is of full rank and has *N*_P_ non-zero eigenvalues. Matrix *K* then has an additional *N*_C_ − *N*_P_ zero eigenvalues. On the other hand, if *s* < 1, 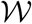 is rank deficient and has *N*_C_ non-zero eigenvalues. Matrix *K* then has a total of *N*_P_ − *N*_C_ zero eigenvalues.

The main features in the spectrum of 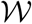 can be appreciated from numerical diagonalisation. For concreteness, we draw the nonzero elements of *M* from a half-normal distribution with probability density function

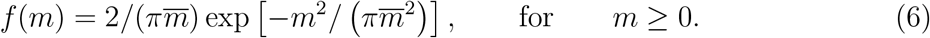

The standard deviation of this distribution is 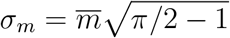. The nonzero elements of *H* are generated in a similar fashion but with using parameter 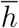. Figure 2 shows the numerical spectrum of instances of matrix 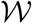 across the range of values of the mutualist ratio *r* with *N*_P_ and *N*_C_ fixed. For *s* > 1 [Fig. 2 (a) and (b)] the spectrum clearly separates into a compact “bulk” spectrum located around *λ* = 0, plus up to two large eigenvalues situated outside the bulk with magnitudes strongly dependent on *r*. The situation for *s* < 1 [Fig. 2 (c) and (d)] is similar, except that here the single bulk is replaced by two disjoint lobes with a further collection of eigenvalues of zero. The large eigenvalues are equally apparent in this case.

**FIG. 2.**
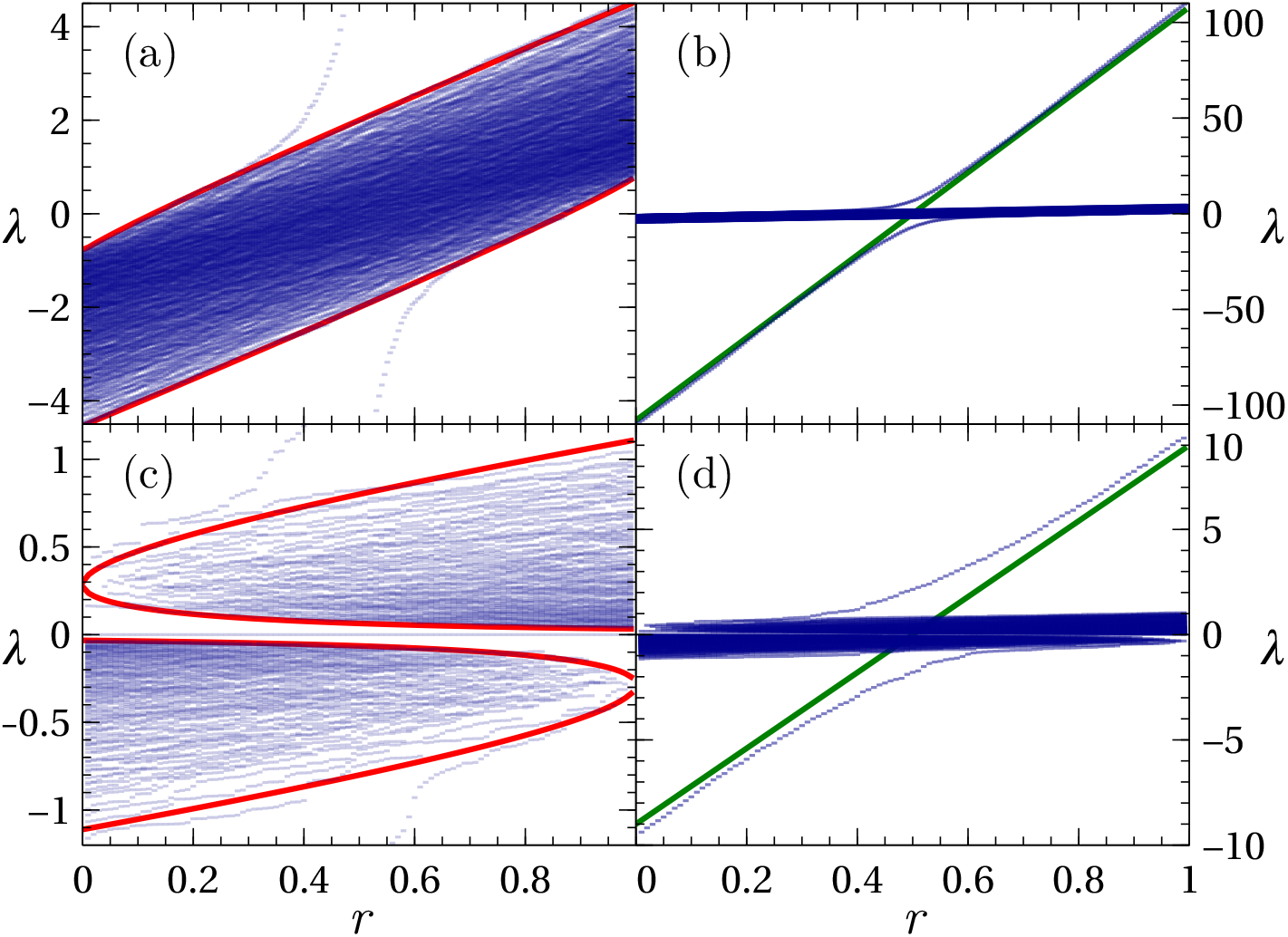
The spectrum of matrix 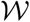 as a function of mutualist ratio *r*. Panels (a) and (b) shows results for a high ratio of consumers to plants, *s* = 6; (c) and (d) show results for a small ratio, *s* = 1/2. The lefthand panels show a close up of the bulk part of the spectrum whilst the righthand panels show the same results on an expanded scale in which large eigenvalues above and below the bulk are readily apparent. Blue symbols are from numerical diagonalisation; red lines on the left are the analytic bulk spectrum edges from Eq. (17) and Eq. (18); green lines on the right are the asymptotic macroscopic value *N*_P_*θ* from Eq. (20). Parameters were *N*_P_ = 200, *c_M_* = *c_H_* = 0.3, *γ* = 1, with non-zero matrix elements chosen from half-normal distributions with 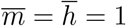.

## IV. EIGENVALUE PHASE TRANSITION

We now analyse the spectrum of 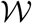 analytically and show that the largest eigenvalue undergoes an eigenvalue phase transition in the *N*_P_ → ∞ limit with the ratios *s* and *r* held fixed. We begin by writing 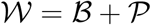 with

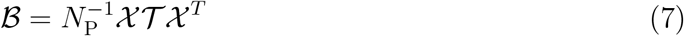

where matrix 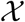 has dimensions *N*_P_ × *N*_C_ and elements

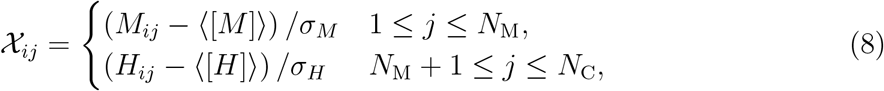

and with 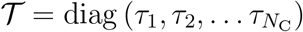 with

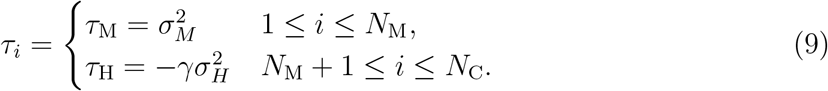

The second matrix in the decomposition of 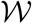 reads

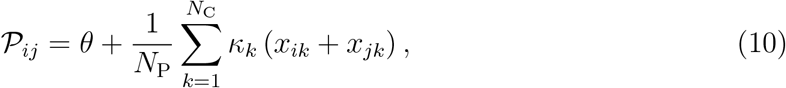

with

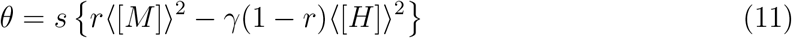

and

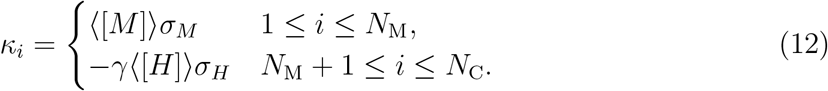

The elements of matrix 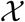 are independent random variables each with zero mean and unit variance. Moments beyond the second play no role in our results in the large-*N*_P_ limit, and thus we take all 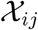 to be distributed identically. Turning to Eq. (10), we observe that the second term is a summation over a large number of independent variables, and thus gives a contribution of the order 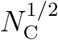 for large *N*_C_. Overall, then, this term scales like 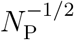 and is therefore negligible in comparison with the first term which scales like *θ* ~ 1. Thus, for 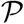, we obtain the approximate rank-1 form

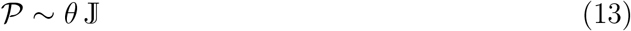

with 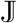 the *N*_P_ × *N*_P_ matrix of ones.

We first consider the matrix 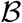 by itself. Let its ordered eigenvalues be 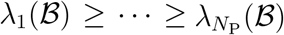 and define the probability measure 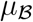 as the limiting empirical eigenvalue distribution 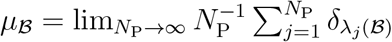. From Ref. 29 [see also Ref. 34], the Stieltjes transform of 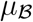 is given by

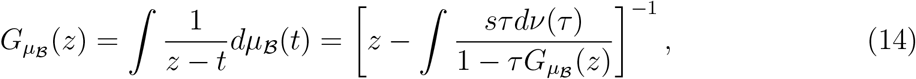

where *ν*(*τ*) is the *N*_P_ → ∞ probability distribution function of the values {*τ*_1_,…, *τ*_*N*_C__}. For the case in hand, this reads

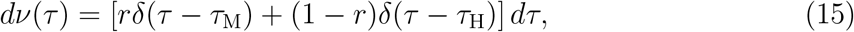

and thus we obtain

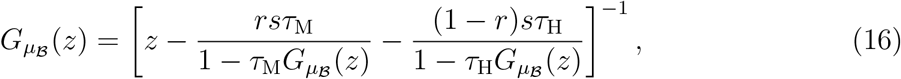

and its inverse

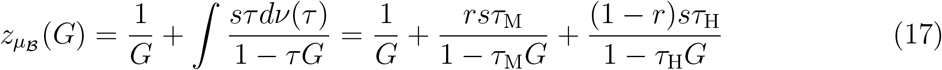

Again from Ref. 29, the edges of connected components of spectrum can be obtained as 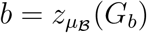 where *G_b_* is a solution of

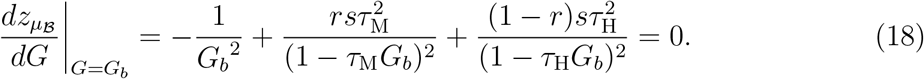

In particular, this gives us *b*_+_ and *b*_−_ the upper and lower edges of the bulk part of the spectrum, i.e. the supremum and infimum of the support of 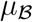. Figure 2 shows the boundaries obtained from Eq. (18) superimposed on the numerical spectra. The boundaries, including *b*_±_, give good approximation to the numerical bulk edges across the entire range of *r*.

Now consider the complete matrix, 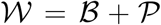. Applying Theorem 2.1 of Ref. 30 to the rank-1 perturbation of Eq. (13), we have that in the *N*_P_ → ∞ limit the uppermost eigenvalue of 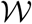 almost surely converges as

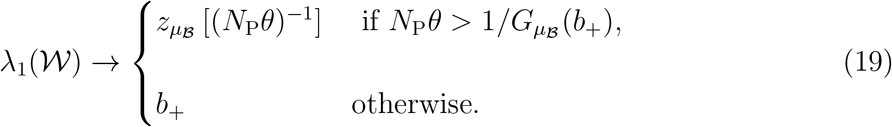

Given that 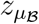 is a monotonically decreasing function at the spectrum edge and that *N*_P_ is large, this becomes

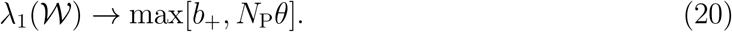

Thus, in the *N*_P_ → ∞ limit, the largest eigenvalue of 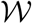 undergoes an abrupt eigenvalue phase transition [31] from a value *b*_+_ ~ (*N*_P_)^0^ below the transition to a value *λ*_1_ = *N*_P_*θ* that is “macroscopic” in the sense that it scales with the size of the ecosystem. This transition occurs at *N*_P_*θ* = *b*_+_ but given the different scaling of the two sides of this equation, the transition point is given by *θ* = 0 in the asymptotic limit. Thus, we see that the phase transition occurs when *r* = *r*_⋆_ with the critical mutualist ratio

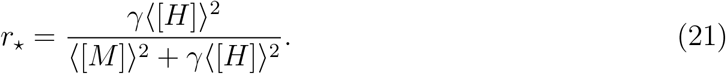

The significance of this is that eigenvalue 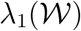 determines the stability of community matrix *K* and is thus an order parameter for a phase transition in the stability of the model. We note that the quantity *N*_P_*θ* is the row sum of matrix 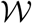, as previously discussed by Ref. 25 in the context of isolated eigenvalues of mutualist matrices. Considering the lowest eigenvalue, we similarly find 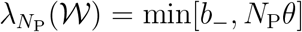. This also exhibits a phase transition but this is of no significance from a point of view of the stability.

Figure 2 shows the macroscopic value *N*_P_*θ* superimposed on the numerical spectra, where it is seen to be close to numerically-obtained large eigenvalues when they lie significantly outside the bulk. Deviations from this good agreement occur in the region close to where *N*_P_*θ* and *b*_±_ cross, and this is a consequence of the finite value of *N*_P_ in this figure.

Finally we note that there is a secondary, less dramatic stability transition implicit in the above results and visible in Fig. 2. This transition occurs at the point *b*_+_ = 0. Below this point, the spectrum of 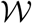 is completely negative [see for example the lefthand edge of Fig. 2 (a)] and thus 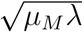 will be imaginary for all eigenvalues. The stability of community matrix *K* will, from Eq. (3), then be given purely by the intraspecific competition term. Above this point, 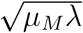 will be real for some *λ* and this will then start to reduce the stability.

## V. FINITE-SIZE EXPRESSION FOR THE LARGE EIGENVALUE

Whilst the above captures the emergence of a macroscopic eigenvalue, Fig. 2 shows that at finite NP the uppermost eigenvalue goes smoothly over from the bulk edge at low *r* to the macroscopic disjoint value at high *r*. In this section we derive an expression for *λ*_1_ that captures this behaviour at large but finite *N*_P_. In doing so, we consider a slightly more general model, namely

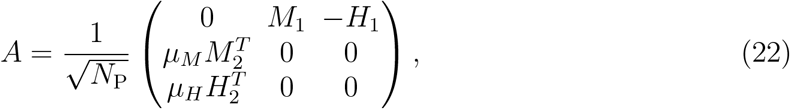

where *M*_1_ and *M*_2_ are random blocks, with relationship left unspecified, and similarly for *H*_1_ and *H*_2_. This allows us to discuss the effect of disorder in the next section. Clearly the model of the previous section is recovered by setting *M*_1_ = *M*_2_ = *M* and *H*_1_ = *H*_2_ = *H*.

With the interaction matrix of Eq. (22), the *N*_P_ × *N*_P_ matrix equivalent of Eq. (5) is

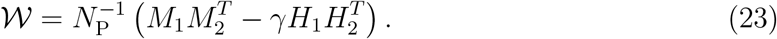

An approximate account of the spectrum of 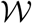 can be obtained by considering this matrix in the eigenbasis of its ensemble average 〈*W*〉. Doing so allows us to identify the scaling properties of different parts of the matrix and derive an approximate expression valid for *N*_P_ ≫ 1. Details of this calculation are described in the Appendix with the results as follows. The spectrum of 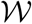 is seen to approximately contain *N*_P_ − 2 eigenvalues at

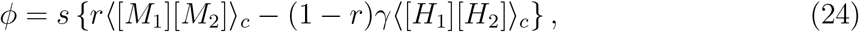

in which 〈[*M*_1_][*M*_2_]〉_*c*_ is the covariance between the matrix elements of *M*_1_ and *M*_2_ (and similarly for 〈[*H*_1_][*H*_2_]〉_*c*_). This set of eigenvalues is the approximate representation of the bulk in this calculation. More importantly, we also obtain two non-trivial eigenvalues given by

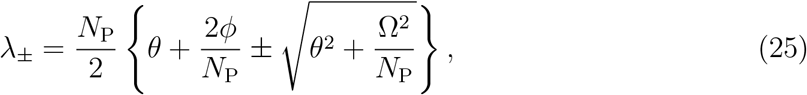

in which, similar to Eq. (11), we have

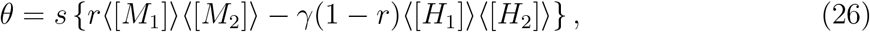

and where

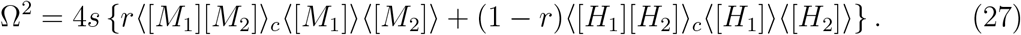

In the asymptotic limit, Eq. (25) gives *λ*_+_ → *N*_P_*θ*, which recovers the macroscopic eigenvalues obtained previously, and *λ*_−_ → *ϕ* which then becomes part of the bulk.

Figure 3 compares the analytic expression *λ*_+_ with the maximum eigenvalue extracted from numerical diagonalisation of 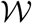 with *M*_1_ = *M*_2_ = *M* and *H*_1_ = *H*_2_ = *H* as in Fig. 2. Results for a range of different values of plant number *N*_P_ are shown. Clear agreement between numerical and analytic results is observed, with the degree of agreement increasing with NP as expected. The only significant deviation between the two at large *N*_P_ occurs at very small values of *r*, where *λ*_+_ overestimates *λ*_1_. In this regime the scaling arguments leading to Eq. (25) break down.

**FIG. 3.**
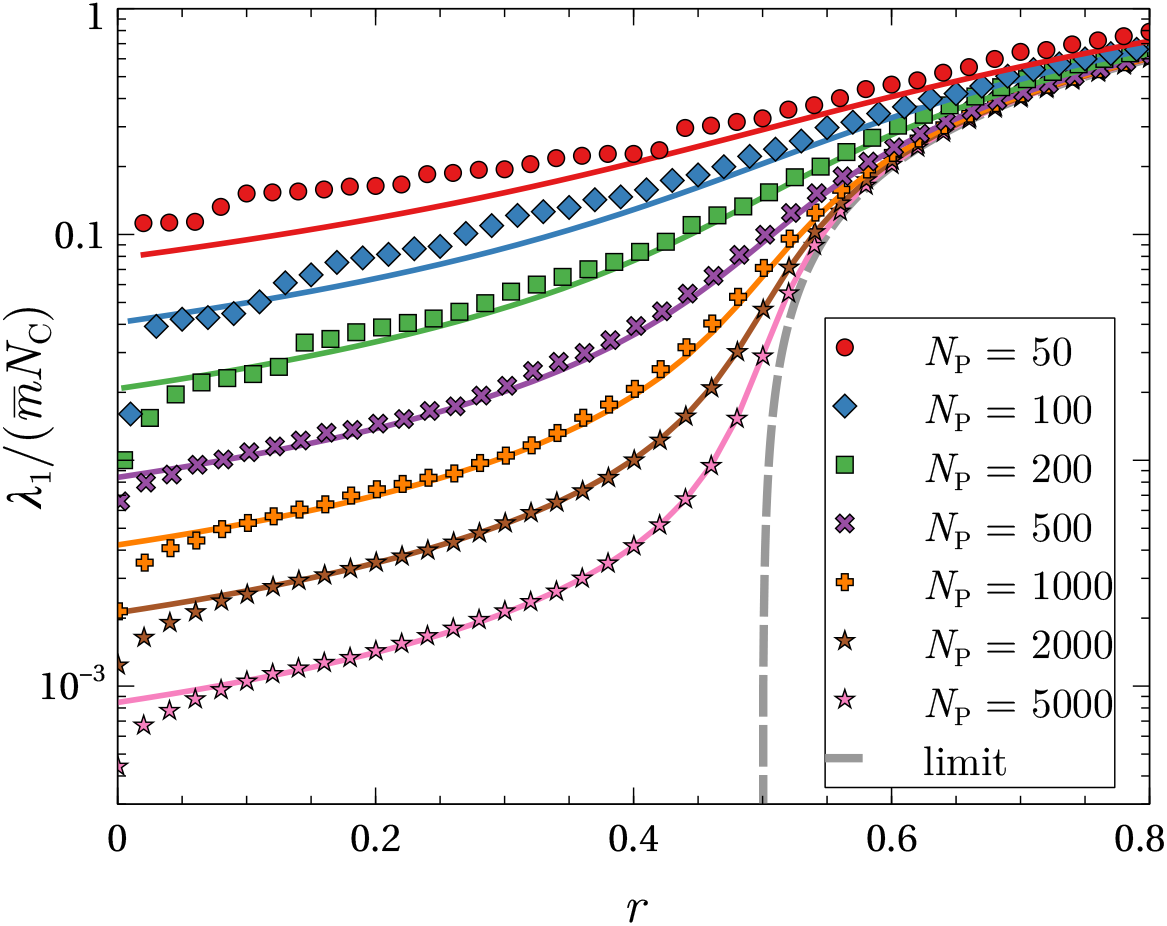
Largest eigenvalue *λ*_1_ of matrix 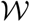 as a function of the mutualist ratio *r* for several values of the plant number *N*_P_ and fixed *s* = *N*_C_/*N*_P_ = 1/2. Symbols show numerical results for a single random instances of W; solid lines show the analytic result *λ*_+_ of Eq. (25); the dashed line shows the asymptotic result of Eq. (20). Other parameters as in Fig. 2.

One additional feature that is revealed by this finite-size analysis is the role of correlations between the strengths of the two directions of the interactions [35]. The influence of these is manifest by the presence of the covariances 〈[*M*_1_][*M*_2_]〉_*c*_ and 〈[*H*_1_][*H*_2_]〉_*c*_ in 〈Ω^2^〉. Since this term enters the expression for *λ*_±_ with a 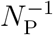 forefactor, correlations can only play a role at finite *N*_P_. In the maximally correlated case, we recover the model of Sec. II for which 〈[*M*_1_][*M*_2_]〉_*c*_ = 〈[*M*]^2^〉_*c*_ as in Eq. (2) and and similarly for *H*. In a scenario where the interaction strengths are completely uncorrelated, we have 〈[*M*_1_][*M*_2_]〉_*c*_ = 〈[*H*_1_][*H*_2_]〉_*c*_ = 0 and 〈Ω^2^〉 vanishes. The prediction for *λ*_1_ in this case becomes the piecewise linear function *λ*_1_ = max [*ϕ*, *ϕ* + *N*_P_*θ*], similar to that found in Sec. IV in the *N*_P_ → ∞ limit. Thus the “avoided crossing” that occurs in the correlated case at large but finite *N*_P_ gives way to an actual crossing when the degree of correlation is zero.

## VI. INTERACTION DISORDER

As defined by their interactions with plants, the consumers in Fig. 1 are either 100% mutualist or 100% herbivore. In this section we look what happens when we move away from this perfectly ordered scenario and swap the types of a random selection of the species interactions. We consider the same tripartite network as before, but with a probability *f*^M^ we swap the (++) signs of the mutualistic interactions to the (+−) signs of an antagonistic one. For the antagonistic interactions, we do the opposite with a probability *f*^H^. The end result is that the animal species no longer act with well defined roles, but differently across their connected plant species. Mathematically, this is incorporated into the framework of Sec. V by selecting the matrix elements of Eq. (22) to be

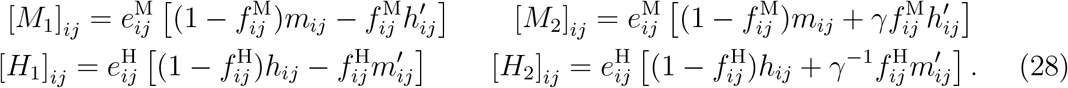

Here, *m_ij_* and *h_ij_* are random variables as before, 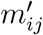 and 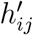 are further independent random variables chosen from the same distributions, and 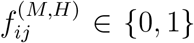 are an additional set of binary random variables that describe whether interaction *ij* in block (*M, H*) is flipped or not. These latter are set with probability 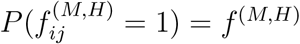.

Using the results of the previous section, we see that, in the asymptotic limit, the largest eigenvalue of the matrix 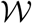 is *λ*_1_ = max[0, *N*_P_*θ*] with *θ* of Eq. (26) in terms of the ensemble averages

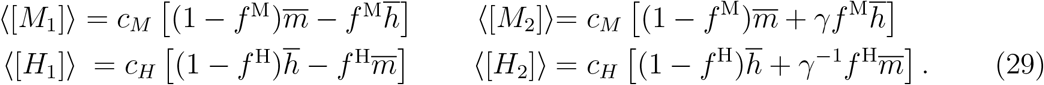

Results are shown in Fig. 4 as phase diagrams in the plane defined by the mutualist ratio *r* and the fraction of swapped interactions *f*^M^ = *f*^H^ = *f*.

**FIG. 4.**
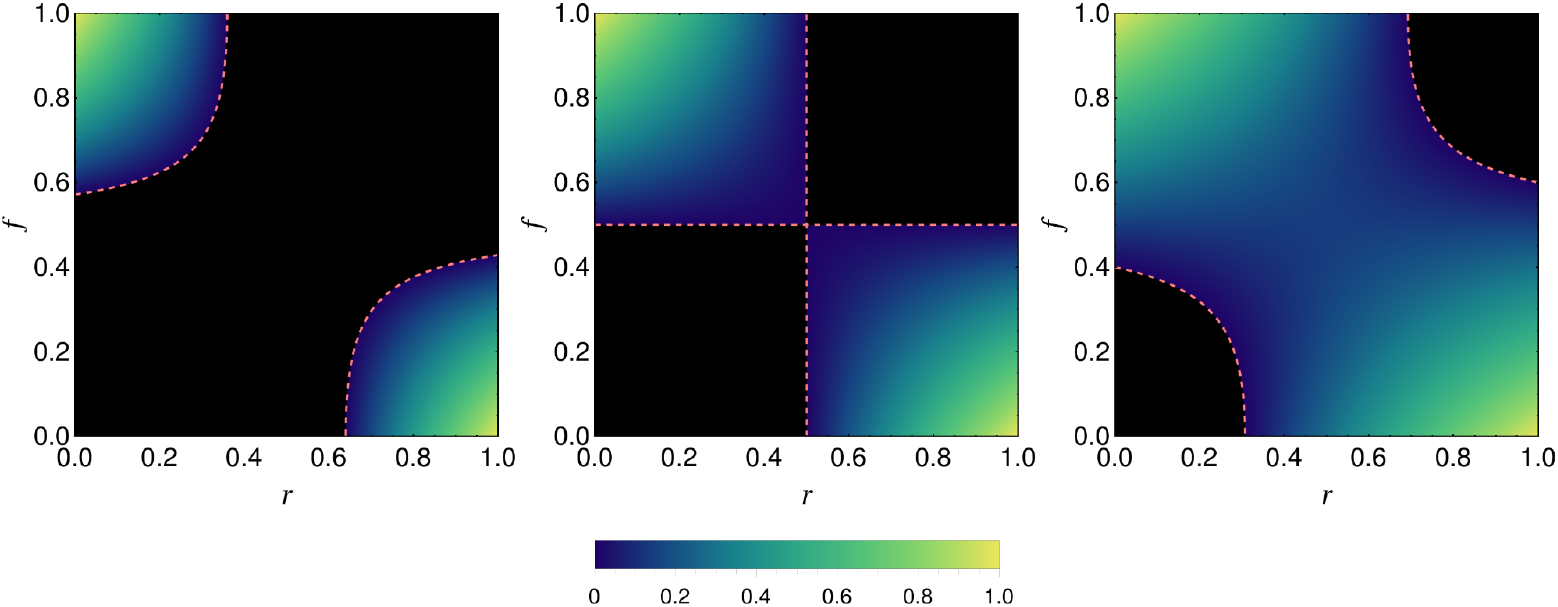
Phase diagrams for the tripartite network of Fig. 1 in which a random fraction *f* of interactions have their type (mutualistic or antagonsitic) switched. Shown is the asymptotic order-parameter eigenvalue *λ*_1_ = max[0, *N*_P_*θ*] with *θ* of Eq. (26) evaluated with the mean values of Eq. (29) as a function of *f* and the mutualist ratio *r*. We have scaled *λ*_1_ by its maximum value (obtained at *r* = *γ* = 1 and *f* = 0) and this removes the dependence on *N*_P_, *s*, and the connectance when *c_M_* = *c_H_* as here. Results are shown for three values of the mutualist interaction strength: 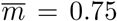, 1, 1.5 from left to right with 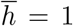 and *γ* = 1. The black regions correspond to the antagonistic phase with *λ*_1_ = 0 and the coloured regions, the mutualistic phase *λ*_1_ > 0. The dashed line represents the phase boundary.

When there is exact symmetry between the interactions strength, i.e. 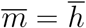 and *γ* = 1 (Fig. 4 middle), the behaviour observed in the fully coherent case, *f* = 0, is preserved at finite *f*, up to a fraction *f* = 1/2. The transition still occurs at *r* = 1/2 and the only change is that the size of order parameter in mutualist phase is diminished. At the point *f* = 1/2 the dominant roles of the “antagonistic” and “mutualistic” species swap and the two phases reverse accordingly. Away from this exact symmetry we have more complicated behaviour. For 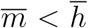 (assuming *γ* = 1) the mutualist phase occupies a diminished area of the phase diagram. For values of *f* close to 1/2 the system remains in the antagonist phase across the whole range of *r* and no instability transition takes place. In contrast, for 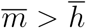, it is the antagonist phase that is diminished and for *f* close to 1/2 the system remains in the mutualist phase for all *r*. Indeed, specifying for simplicity the case of *c_M_* = *c_H_*, *f*^M^ = *f*^H^ = *f* and *γ* = 1 (as in Fig. 4), we find the value of *r* for which *θ* = 0 to be

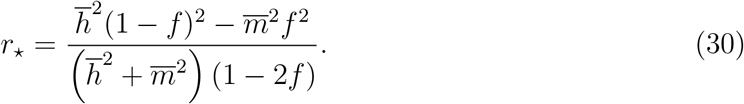

For a phase transition to occur, we require that 0 < *r*_⋆_ < 1, which gives

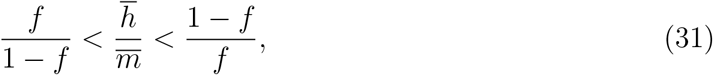

as the condition on the interaction and disorder strengths for the existence of a phase transition.

## VII. DISCUSSION

We have studied the stability of a community matrix with random elements organised according to the tripartite structure of Fig. 1. This structure can be viewed as the mergence of two bipartite networks with changes in the mutualist ratio *r* interpolating between them. At *r* = 0 we have a bipartite predator-prey model, whose community matrix has purely imaginary eigenvalues [32]; at *r* = 1 we have a bipartite mutualist model, whose stability is determined by a single large real eigenvalue [25]. We have shown here that the emergence of this macroscopic eigenvalue as a function of *r* is abrupt in the asymptotic limit, and that this represents an eigenvalue phase transition that takes place when *r* reaches the critical value *r*_⋆_. The behaviour of the model is thus split into two distinct regimes. For *r* < *r*_⋆_, we obtain an “antagonistic phase” in which the community matrix *K* can be stabilised by a small (i.e. order 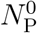) value of the intraspecific competition *d*. In this regime, changing the number of mutualists by a small amount does not appreciably affect the stability of the system. In contrast, for *r* > *r*_⋆_ we obtain a “mutualist phase” in which stability is governed by the macroscopic eigenvalue such that the community matrix requires a large intraspecific competition, 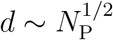, to stabilise it. Moreover, in this phase, small changes in the mutualist number lead to large changes in *λ*_1_ and hence dramatic changes in the stability of the system. Whilst we have framed this discussion in terms of the behaviour as a function of the ratio *r*, the phase transition can also be driven by changes in other parameters, e.g. mean interaction strengths or connectance.

A large positive eigenvalue of the community matrix is associated with the growth of populations away from the equilibrium values. The rapidity seen here is a consequence of the positive feedback of “mutual benefaction” associated with the mutualistic interactions [36]. It should be born in mind, however, that the community matrix represents a linearisation of a more complex/detailed dynamical model. Thus, instability should not be interpreted as an uncontrolled growth, but rather as the indication of the shift of the ecosystem away from its equilibrium to a different one, the properties of which are outside the scope of the original model. This new equilibrium may well include fewer species than in the starting community.

The scaling in Eq. (1) is chosen such that the bulk spectrum of *K* converges in the *N*_P_ → ∞ limit. This follows from the convergence of the spectrum of 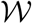 with bulk edges *b*_±_ ~ (*N*_P_)^0^. In the spectrum of *K*, this leaves the scaling of the macroscopic eigenvalue as 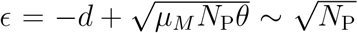. An slightly different choice would be to replace the 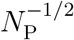 forefactor out the front of Eq. (1) with 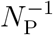. This would make extreme eigenvalue of *K* converge to 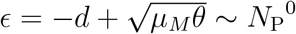 in the limit, whilst the bulk would shrink to a point at *ϵ* = −*d*. Given the different scalings of the two parts of the spectrum, we might also consider a slightly different model in which this forefactor in Eq. (1) is omitted and we instead choose to scale the matrix-element distribution such that 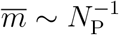 and 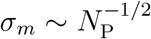 and similarly for the herbivores. This situation would then be similar to the scaling in e.g. Ref. 37 and 38. This choice invalidates the assumptions upon which above calculations are based (operator 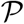 can no longer be approximated as above; the scaling arguments of Sec. V no longer hold). However, because here (and unlike references just cited) the matrix elements are strictly non-negative, *m_ij_, h_ij_* ≥ 0, the only way in which to obtain the required scaling of 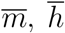, *σ_m_* and *σ_h_* is with a distribution that is concentrated at values of *m* extremely close to zero but which nevertheless has a very long tail. This scaling therefore converts most of the interactions in the community matrix into vanishing small ones (with a few very large ones to achieve the required mean) which seems rather at odds with the set-up of the model.

Phase transitions of the type described here have been described in related ecological models. Ref. 39 discussed an *S* × *S* community matrix with random elements distributed homogeneously about a finite mean, and showed that this can give rise to a macroscopic eigenvalue of size 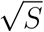 (in the scaling convention pursued here). This model was also considered by Ref. 35 and Ref. 40 who described the emergence of this macroscopic eigenvalue as a phase transition from stability to instability as the mean value of the matrix elements is increased. The connection with the antagonist-mutualist system discussed here is that an increased mean interaction strength could arise from an increase in the number of mutualistic interactions. This scenario was discussed by Ref. 27 in another homogeneous model, introduced by Ref. 26, that consists only of antagonists and mutualists. In these homogeneous models, the macroscopic eigenvalue can be understood in terms of the row sum of the community matrix *K* [25]. In contrast, in the structured model discussed here, it is the row sum of the folded matrix 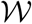 that is important.

Ref. 13 considered the same tripartite topology as discussed here. Their conclusion was that stability was enhanced by the connectance and species diversity of the mutualistic subnetwork but decreased by the connectance and diversity of the antagonistic one. This is opposite to the behaviour described here (where at best increased mutualistic interactions can leave the stability unaltered in the strict asymptotic limit). There are several reasons for this discrepancy. Ref. 13 consider a relatively small number of species (compared with the limiting behaviour here) described by a more detailed dynamical model with a particular subset of parameters that guarantees that the mutualist subsystem was independently stable. Perhaps even more importantly is that the stability to which they refer is that of a final equilibrium state, obtained through the time-evolution of the model. This will in general have fewer species than the starting community and possess additional structure. It is also clear that non-random structure can affect the stability of such networks, as Ref. 17 show in their comparison of random and empirical networks.

It seems therefore that an important future direction is to look at the feasibility of dynamical systems with interaction topology similar to that in Fig. 1. This would also allow better connection with the persistence studies of Ref. 13 and functional extinctions of Ref. 28. The dynamical cavity method has proved very useful for studying feasibility in homogeneous models [37, 38], and it remains on open possibility that this approach can be extended to structured models as studied here. Studying related few-species models, Ref. 41 commented that the destablizing effect of mutualisms “… is more than canceled by an increased chance of feasibility”, and it will be interesting to see whether this also applies for large networks of interacting species and, in particular, in the presence of a phase transition.

Finally we note that our analysis can be generalised to a more complex merged networks, reminiscent of that in Ref. 15, which consist of a central guild of *N*_P_ species (plants in Fig. 1) that interacts with a set of *G* other guilds, all of which are non-interacting with one another. Similar to Eq. (22), the interacting part of community matrix will be of the block form

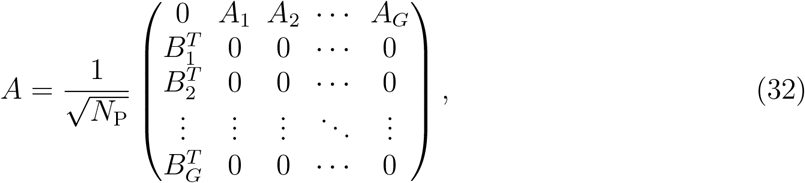

where *A_i_* and *B_i_* are random matrices of dimension *N*_P_ × *N_i_* with *N*_i_ the number of species in guild *i*. In contrast to Eq. (22), the interaction signs [e.g. (+, −) for antagonisms] here are given by the matrix elements, rather than being explicit in the block structure. Similar to above, finding the spectrum of this *K* reduces to the problem of finding the spectrum of 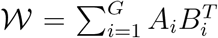. This model possess a maroscopic eigenvalue, 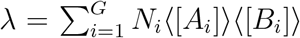 in the asymptotic limit, whose position relative to the bulk around zero determines the stability. From the interaction signs, a guild of mutualists gives a positive contribution to *λ*, whilst and antagonist give a negative one. Competitive interactions in which elements of both *K_i_* and *B_i_* would be both negative would then also give a positive contribution to *λ*, thus moving the system in the direction of instability.

## ACKNOWLEDGMENTS

This work was supported by Newcastle University’s Institute for Sustainability. We acknowledge helpful discussions with Darren M. Evans.

## Appendix A Finite-*N*_P_ expression for the largest eigenvalue

In this appendix we provide details on how the approximate expression for *λ*_1_ in Sec. V is obtained. Our starting point is the more general model of Eq. (22) and the corresponding 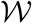 matrix of Eq. (23). The ensemble average of 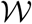 evaluates as

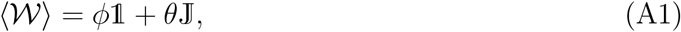

with *ϕ* and *θ* as in Eq. (24) and Eq. (26) and where 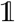 and 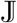 are the *N*_P_ × *N*_P_ unit matrix and matrix of ones, respectively. The eigenvalues of 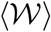 are *ϕ* + *N*_P_*θ* and *ϕ*, this latter being (*N*_P_ − 1)-fold degenerate. The eigenvector *u*^(1)^ belonging to eigenvalue *ϕ* + *N*_P_*θ* has elements

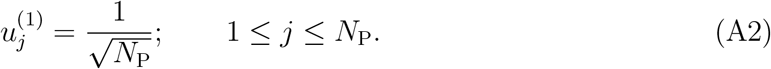

In the degenerate subspace, we choose the eigenvector set (*u*^(*α*)^: 2 ≤ *α* ≤ *N*_P_} with elements

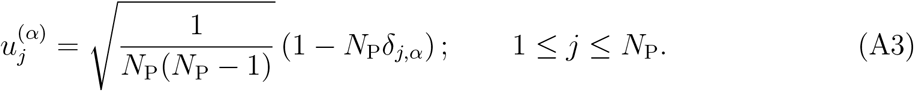

These states are normalised, orthogonal to *u*^(1)^ but not orthogonal to each other. Indeed we have

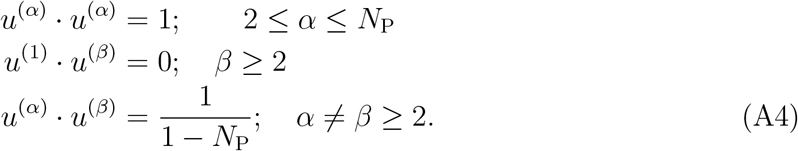

These eigenvectors are easier to work with than orthogonalised ones, and since we work in the *N*_P_ limit, the effect of the finite overlap *u*^(*α*)^ · *u*^(*β*)^ is negligible.

Arranging *u*^(*α*)^ as columns of matrix *U*, the full *W* matrix in this (ensemble-averaged) basis is 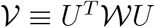, with matrix elements

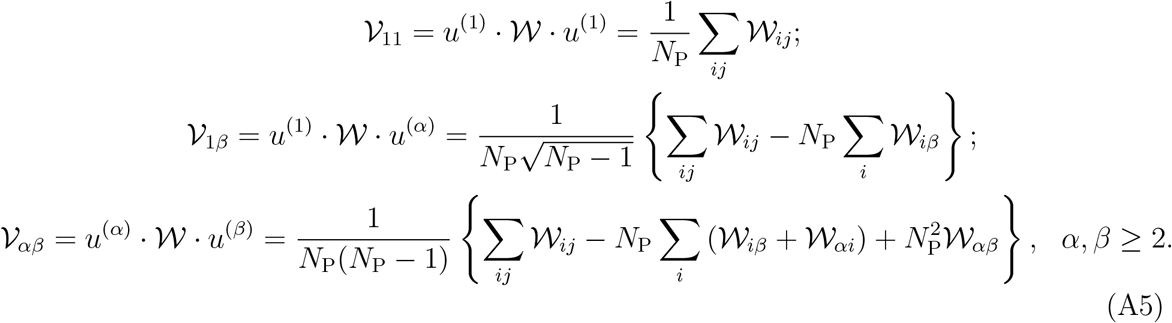

Considering the scaling of the ensemble averages of these quantities, we find 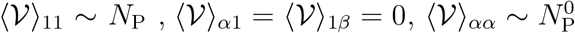, and 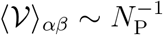 for *α* ≠ *β*. Similarly, the variances scale as 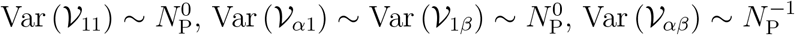. Thus, with *N*_P_ large, we drop 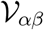 for *α* ≠ *β* as being negligible, and replace the diagonal elements with their ensemble averages, as their fluctuations are negligible in comparison. Due to the vanishing of their mean, the elements 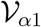 and 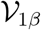 are left as is. Taken together, this gives the approximation

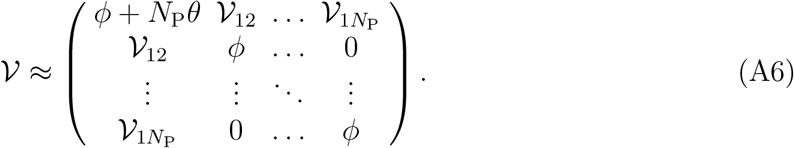

This can be diagonalised exactly. We obtain *N*_P_ − 2 eigenvalues of value *ϕ* and two nontrivial ones. These latter have the form of *λ*_±_ from Eq. (25) with

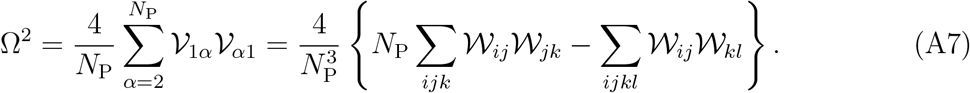

Evaluating the leading term in the ensemble average of Ω^2^ we find

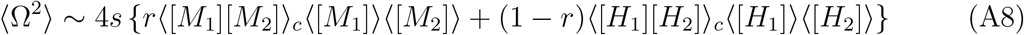

A similar calculation for the variance show that it scales like 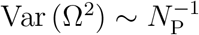 such that the fluctuations about the mean vanish in the limit. We thus approximate Ω^2^ with 〈Ω^2^〉, thus giving the result stated in Eq. (27).

